# C-type lectin-like receptor 2 (CLEC-2)-dependent DC migration is controlled by tetraspanin CD37

**DOI:** 10.1101/227918

**Authors:** Charlotte M de Winde, Alexandra L Matthews, Sjoerd van Deventer, Alie van der Schaaf, Neil D Tomlinson, Erik Jansen, Johannes A Eble, Bernhard Nieswandt, Helen M McGettrick, Carl G Figdor, Sophie E Acton, Michael G Tomlinson, Annemiek B van Spriel

**Author notes:** Authors contributed equally to the work (Alexandra L Matthews, Sjoerd van Deventer, Sophie E Acton, Michael G Tomlinson).

## Abstract

Cell migration is central to evoke a potent immune response. Dendritic cell (DC) migration to lymph nodes is dependent on the interaction of C-type lectin-like receptor 2 (CLEC-2) expressed by DCs, with podoplanin expressed by lymph node stromal cells (LNSCs). However, the underlying molecular mechanisms by which CLEC-2 influences DC migration remain elusive. Here, we show that CLEC-2-dependent DC migration is tightly controlled by tetraspanin CD37, a membrane-organizing protein. Our findings demonstrate a specific molecular interaction between CLEC-2 and CD37. Myeloid cells lacking CD37 (*Cd37-/-*) expressed less CLEC-2 on their surface compared to wild-type cells, indicating that CD37 is required to stabilize membrane expression of CLEC-2. In addition, CLEC-2-expressing DCs lacking CD37 showed impaired adhesion, migration velocity and displacement on LNSCs. Moreover, *Cd37-/-* DCs failed to form actin protrusions in a 3D collagen matrix upon podoplanin-induced CLEC-2 stimulation, phenocopying CLEC-2-deficient DCs (CD11c^ΔCLEC-2^). Microcontact printing experiments revealed that CD37 is required for CLEC-2 recruitment in the membrane to its ligand podoplanin. This study demonstrates that tetraspanin CD37 controls CLEC-2 membrane organization and provides new molecular insights underlying CLEC-2-dependent DC migration.

## Introduction

Cell migration is a key process in the initiation of immune responses (Worbs et al., 2017). Upon encountering a foreign antigen, dendritic cells (DCs) migrate to secondary lymphoid organs (i.e. lymph nodes, spleen) to present antigen on major histocompatibility complex (MHC) and activate T and B lymphocytes. En route, DCs move along podoplanin-expressing lymph node stromal cells (LNSCs), such as lymphatic endothelial cells (LECs) and fibroblastic reticular cells (FRCs). The interaction between C-type lectin-like receptor 2 (CLEC-2) on DCs with podoplanin on LNSCs is essential for optimal DC migration to and within the lymph node (Acton et al., 2012). Despite the important role of CLEC-2 in DC migration, the molecular mechanisms underlying CLEC-2-dependent cell migration remain to be elucidated.

CLEC-2 (encoded by the gene *Clec1b*) is expressed on platelets (Suzuki-Inoue et al., 2006) and myeloid immune cells, such as DCs, macrophages and neutrophils (Colonna et al., 2000; Lowe et al., 2015; Mourão-Sá et al., 2011). CLEC-2 plays a key role in fetal development of the lymphatic vasculature, as demonstrated by *Clec1b*-knockout mice which are embryonically lethal (Bertozzi et al., 2010; Suzuki-Inoue et al., 2010). Besides podoplanin, the snake venom toxin rhodocytin is another ligand for CLEC-2. Both ligands initiate downstream signaling via Syk resulting in cell activation (Fuller et al., 2007; Hughes et al., 2010; Suzuki-Inoue et al., 2006). Intracellular Syk-binding requires dimerization of CLEC-2 receptors, since each CLEC-2 receptor contains only a single tyrosine phosphorylation (YXXL) motif. This makes CLEC-2 a hemITAM (hemi-Immunoreceptor Tyrosine-based Activation Motif) C-type lectin receptor (CLR), similar to its homologous family member Dectin-1 (CLEC7A) that recognizes β-glucans in fungal cell walls (Brown and Gordon, 2001; Fuller et al., 2007).

The organization of receptors in the plasma membrane of DCs plays a pivotal role in immune cell function (Zuidscherwoude et al., 2014; Zuidscherwoude et al., 2017a). For proper ligand binding and initiation of signaling, CLRs are dependent on localization into membrane microdomains, such as lipid rafts and tetraspanin-enriched microdomains (TEMs) (Figdor and van Spriel, 2009; Zuidscherwoude et al., 2014). TEMs, also referred to as the tetraspanin web, are formed by the interaction of tetraspanins, a family of four-transmembrane proteins, with each other and partner proteins (Charrin et al., 2009; Hemler, 2005; Levy and Shoham, 2005; Zimmerman et al., 2016; Zuidscherwoude et al., 2015). As such, TEMs have been implicated in fundamental cell biological functions, including proliferation, adhesion and signaling (Charrin et al., 2009; Hemler, 2005; Levy and Shoham, 2005). Earlier work indicated that CLEC-2 clustering and signaling in blood platelets is dependent on lipid rafts (Manne et al., 2015; Pollitt et al., 2010). CLEC-2 was shown to be present as single molecules or homodimers on resting platelets, and larger clusters were formed upon rhodocytin stimulation (Hughes et al., 2010; Pollitt et al., 2014), but the molecular mechanism underlying ligand-induced CLEC-2 clustering is yet unknown. More is known about the organization of Dectin-1, a CLEC-2 homologous family member, on immune cells. Dectin-1 molecules need to be reorganized into a “phagocytic cup” to bind and phagocytose particulate, and not soluble, β-glucan (Goodridge et al., 2011). Furthermore, Dectin-1 has been proposed to be present in lipid rafts (De Turris et al., 2015; Xu et al., 2009) or tetraspanin (CD63, CD37) microdomains (Mantegazza et al., 2004; Meyer-Wentrup et al., 2007; Yan et al., 2014) on myeloid cells.

Tetraspanin CD37 is exclusively expressed on immune cells with highest expression on B lymphocytes and DCs (de Winde et al., 2015). The importance of CD37 in the immune system has been demonstrated in CD37-deficient mice (*Cd37-/-*) that have defective humoral and cellular immune responses (van Spriel et al., 2004; van Spriel et al., 2012). Interestingly, DCs that lack CD37 showed impaired spreading, adhesion and migration, leading to defective initiation of the cellular immune response (Gartlan et al., 2013; Jones et al., 2016). Since CLEC-2 plays an important role in DC migration (Acton et al., 2012), and its homologous receptor Dectin-1 has been shown to interact with tetraspanin CD37 (Meyer-Wentrup et al., 2007), we hypothesized that CD37 may influence CLEC-2 membrane organization and thereby controls DC migration. In this study, we show that CLEC-2 interacts with CD37. Moreover, CLEC-2-dependent actin protrusion formation by DCs and recruitment of CLEC-2 to podoplanin is dependent on CD37 expression. These results provide evidence that tetraspanin CD37 is required for CLEC-2 recruitment in the plasma membrane in response to podoplanin, and as such plays an important role in CLEC-2-dependent DC migration.

## Results

### CLEC-2 interacts with tetraspanin CD37

To investigate whether CLEC-2 interacts with tetraspanins, we performed co-immunoprecipitation experiments with HEK-293T cells transfected with human CLEC-2-MYC with or without a range of FLAG-tagged human tetraspanin constructs. These experiments revealed that CLEC-2 interacts with CD37, but not with four other tetraspanins used as controls (CD9, CD63, CD151 or CD81) under conditions using 1% digitonin (Figure 1A-B). Interactions preserved in 1% digitonin are associated with primary (direct) interactions between a tetraspanin and its partner proteins (Serru et al., 1999). The strength of the interaction between CD37 and CLEC-2 was comparable with two well-established primary interactions between tetraspanins and their partner proteins: CD9 with CD9P1 (Charrin et al., 2001) and Tspan14 with ADAM10 (Dornier et al., 2012; Haining et al., 2012) (Figure 1C-D). Thus, these experiments show that CLEC-2 specifically interacts with tetraspanin CD37.

**Figure 1.**
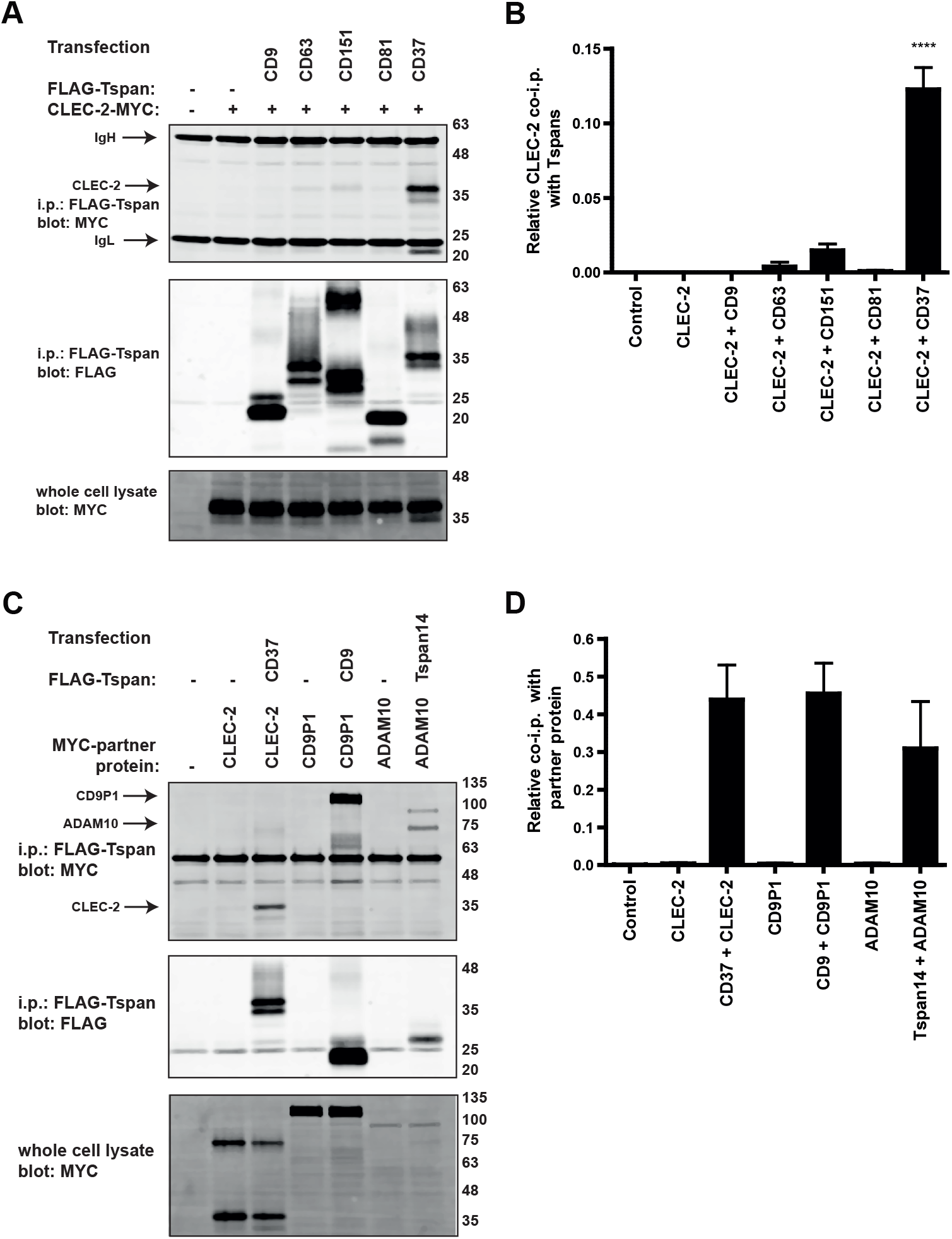
CLEC-2 specifically interacts with tetraspanin CD37. **(A)** HEK-293T cells were co-transfected with MYC-tagged human CLEC-2 and FLAG-tagged human tetraspanins (CD9, CD63, CD151, CD81 or CD37), or mock transfected (-). Cells were lysed in 1% digitonin and immunoprecipitated with an anti-FLAG antibody. Immunoprecipitated proteins were blotted with anti-MYC antibody (top panel) or anti-FLAG antibody (middle panel). Whole cell lysates were probed with the anti-MYC antibody (bottom panel). **(B)** Quantification of (A); amount of MYC-tagged CLEC-2 co-precipitated was normalized to the amount of tetraspanins on the beads. Data are shown as mean+SEM from three independent experiments. Data were normalized by log transformation and statistically analyzed using one-way ANOVA with a Tukey’s multiple comparison test compared with the mean of every other column (****p<0.0001). **(C)** HEK-293T cells were co-transfected with MYC-tagged human CLEC-2, CD9-P1 or ADAM10 expression constructs and FLAG-tagged CD37, CD9 or Tspan14 tetraspanins, or mock transfected (-). Cells were lysed in 1% digitonin and immunoprecipitated with an anti-FLAG antibody. Immunoprecipitated proteins were blotted with anti-MYC antibody (top panel) or anti-FLAG antibody (lower panel). Whole cell lysates were probed with the anti-MYC antibody (middle panel). **(D)** Quantification of (C); amount of MYC-tagged partner co-precipitated was normalized to the amount of tetraspanins on the beads. Data are shown as mean+SEM from three independent experiments.

### CD37-deficient myeloid cells show decreased CLEC-2 surface expression and increased CLEC-2-dependent IL-6 production

We investigated CLEC-2 membrane expression on immune cells of *Cd37-/-* mice. Naïve *Cd37-/-* myeloid cells expressed significantly lower CLEC-2 levels compared to *Cd37+/+* (WT) splenocytes (Figure 2A). It was reported that CLEC-2 expression was increased on myeloid cells upon *in vivo* LPS stimulation (Lowe et al., 2015; Mourão-Sá et al., 2011). Therefore, we analyzed CLEC-2 expression on different immune cell subsets from spleens of *Cd37-/-* and WT mice that were stimulated with LPS *in vivo.* CLEC-2 membrane expression was substantially increased by LPS, but this increase was significantly lower on LPS-stimulated *Cd37-/-* myeloid cells (DCs, macrophages, granulocytes), compared to WT myeloid cells (Figure 2B-C). This was in contrast to LPS-stimulated WT and *Cd37-/-* lymphoid cells (B cells, T cells and NK cells) that expressed comparable CLEC-2 levels (Figure 2C). As a control, we investigated CLEC-2 expression on mouse platelets, which do not express CD37 (Zeiler et al., 2014), and observed similar CLEC-2 expression on platelets of WT and *Cd37-/-* mice (Supplementary Figure 1).

**Figure 2.**
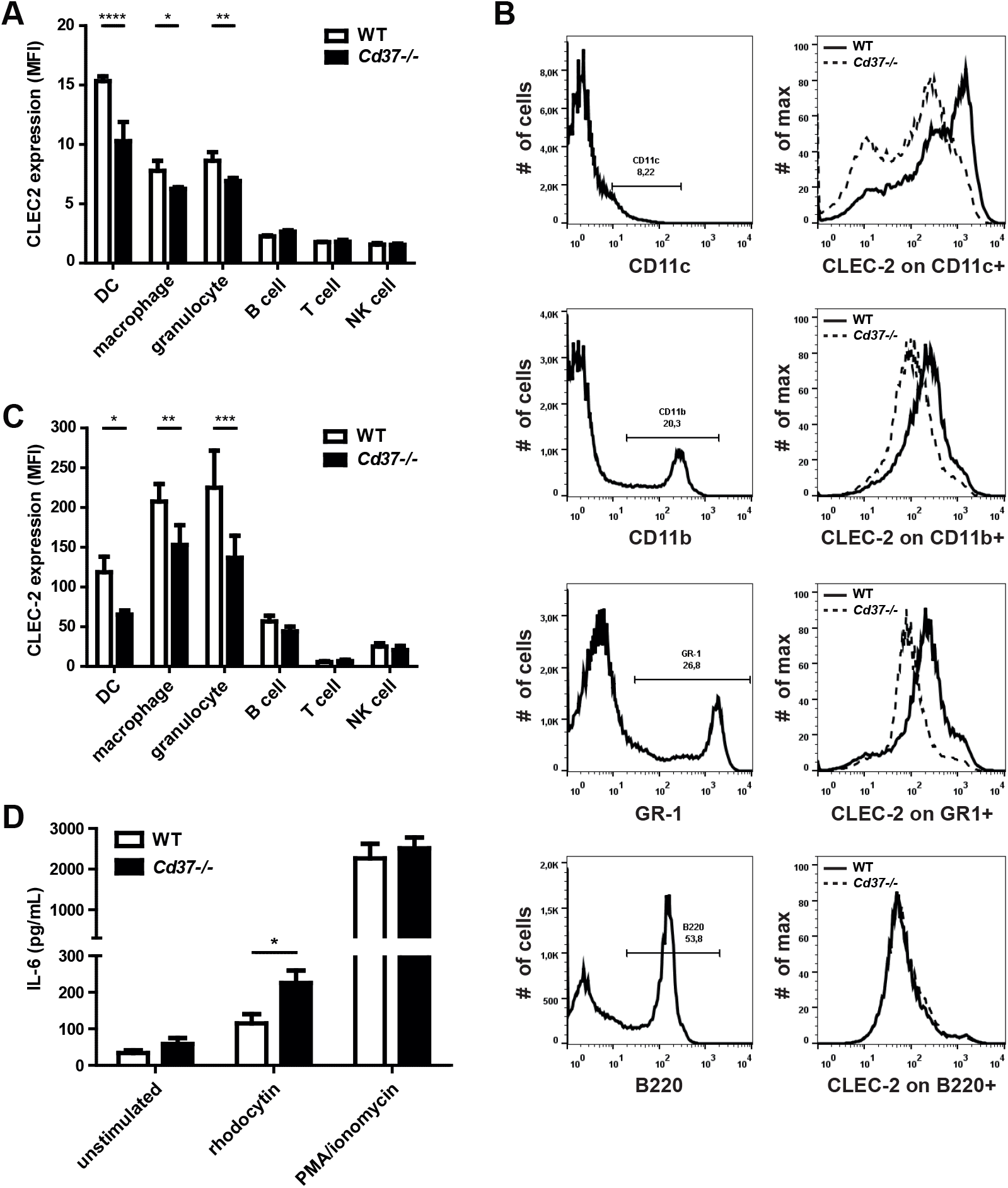
CD37-deficiency impairs CLEC-2 expression and myeloid cell function. **(A-C)** CD11c was used to stain DCs, CD11b for macrophages, GR-1^high^ for granulocytes, B220 for B cells, CD3ε for T cells and NK1.1 for NK cells. **(A)** Quantification of flow cytometric analysis of CLEC-2 expression of naïve WT (white bars) and *Cd37-/-* (black bars) cells. Values are corrected for isotype controls. MFI = mean fluorescence intensity. Data are shown as mean+SD from n=2-3 mice per genotype. Two-way ANOVA with Sidak’s multiple comparisons test, *p<0.05, **p<0.01, ****p<0.0001. **(B)** Flow cytometric analysis of CLEC-2 expression (antibody clone INU-1) on splenic immune cell subsets in WT (black line) and *Cd37-/-* (dashed line) mice 24h post-i.p. injection with LPS. Representative FACS plots for one WT and one *Cd37-/-* mouse per immune cell type are shown. **(C)** Quantification of flow cytometric analysis of CLEC-2 expression of *in vivo* LPS stimulated WT (white bars) and *Cd37-/-* (black bars) cells. Values are corrected for isotype controls. MFI = mean fluorescence intensity. Data are shown as mean+SD from n=2-3 mice per genotype. Two-way ANOVA with Sidak’s multiple comparisons test, *p<0.05, **p<0.01, ***p<0.001. **(D)** IL-6 production (in pg/mL) by total splenocytes from naive WT (white bars) and *Cd37-/-* (black bars) mice after *ex vivo* stimulation with medium (unstimulated, negative control), 15μg/mL rhodocytin (CLEC-2 agonist) or PMA/ionomycin (positive control). Data are shown as mean+SEM from 3 independent experiments, total n=6-8 mice per genotype. Non-parametric Mann-Whitney test, two-tailed, *p=0.0215.

Next, we investigated whether decreased CLEC-2 expression on *Cd37-/-* immune cells had functional consequences by analyzing cytokine production. *Cd37-/-* splenocytes produced significantly more interleukin-6 (IL-6) compared to WT splenocytes upon stimulation with the CLEC-2 ligand rhodocytin (Figure 2D). This was not due to a general defect of the *Cd37-/-* immune cells since stimulating these cells with PMA/ionomycin resulted in equivalent IL-6 production compared to WT cells (Figure 2D). Thus, presence of CD37 is important for regulation of CLEC-2 membrane expression on myeloid cells and CD37 inhibits CLEC-2-dependent cytokine production.

### CD37 controls DC migration and protrusion formation in response to podoplanin

To investigate whether CD37 was required for the migratory capacity of CLEC-2-expressing (CLEC-2+) DCs, we performed static adhesion and migration assays of WT and *Cd37-/-* CLEC-2+ (LPS-stimulated) DCs on LECs (Figure 3A). Adhesion of both WT and *Cd37-/-* CLEC-2+ DCs to inflamed (TNFα-stimulated) LECs was stable for the duration of the experiment. Interestingly, the percentage of *Cd37-/-* DCs adhering to the inflamed LECs was reduced when compared to WT DCs (Figure 3B). Moreover, migration velocity and mean square displacement of DCs were significantly decreased in absence of CD37 (Figure 3C-D, Supplementary movies 1A-B).

**Figure 3.**
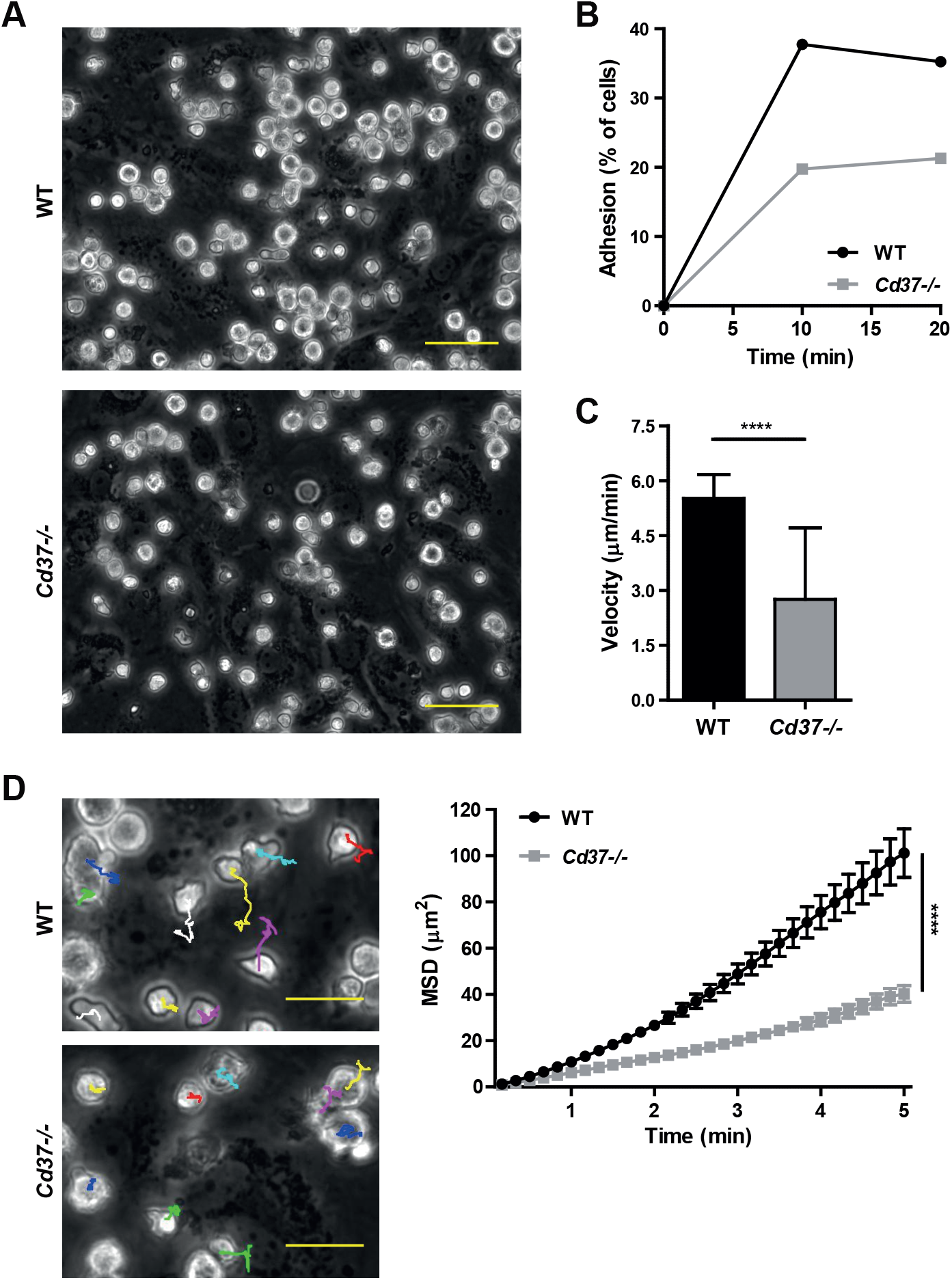
CD37 controls DC adhesion, migration velocity and displacement on lymphatic endothelial cells. **(A)** Adherence of WT (left) and *Cd37-/-* (right) DCs on TNFα-stimulated lymphatic endothelial cells (LEC). One representative image per genotype is shown. Scale bar = 50μm. **(B)** Adhesion (% of total cells added per well) of WT (black line) and *Cd37-/-* (grey line) DCs on LECs for the duration of the experiment. Data shown from one representative experiment. Experiments were repeated two times yielding similar results. **(C)** Migration velocity (μm/min) of WT (white box) and *Cd37-/-* (grey box) DCs over LECs. Bars represent median with interquartile range from two independent experiments, n=107-115 cells per genotype, respectively. Non-parametric Mann Whitney test, two-tailed, ****p<0.0001. **(D)** Left: zoom of field of view shown in **(A)** with individual cell tracks. Tracking paths of each cell of one representative experiment are shown in Supplementary Movies 1A-B. Upper image= WT, lower image = *Cd37-/-.* Scale bar = 25μm. Right: Mean square displacement (MSD, in μm^2^) of WT (black line) and *Cd37-/-* (grey line) DCs on LEC. Data are shown as mean±SEM from two independent experiments. Two-way ANOVA with Sidak’s multiple comparisons, ****p<0.0001 at t=5 min.

To investigate whether cell morphological changes underlying DC migration are CLEC-2-dependent, we analyzed the ability of DCs to form protrusions in response to the CLEC-2 ligand podoplanin in a 3D collagen matrix. In response to podoplanin, the protrusion length and morphology index of WT DCs was significantly increased compared to unstimulated cells, as previously described (Acton et al., 2012) (Figure 4A,C-D and Supplementary Figure 2). DCs lacking CD37 were capable of forming short actin protrusions (Figure 4A-B, Supplementary Figure 2). However, *Cd37-/-* DCs, despite expressing similar levels of CLEC-2 (data not shown), were unable to increase protrusion length and morphology index in response to podoplanin, instead phenocopying the defect seen in DCs lacking CLEC-2 (CD11c^ΔCLEC-2^; (Acton et al., 2012; Acton et al., 2014)) (Figure 4A,C-D). Together, these results demonstrate that CD37-deficiency, similar to CLEC-2-deficiency, results in aberrant DC adhesion and migration on LNSCs, and decreased actin protrusion formation in response to the CLEC-2 ligand podoplanin.

**Figure 4.**
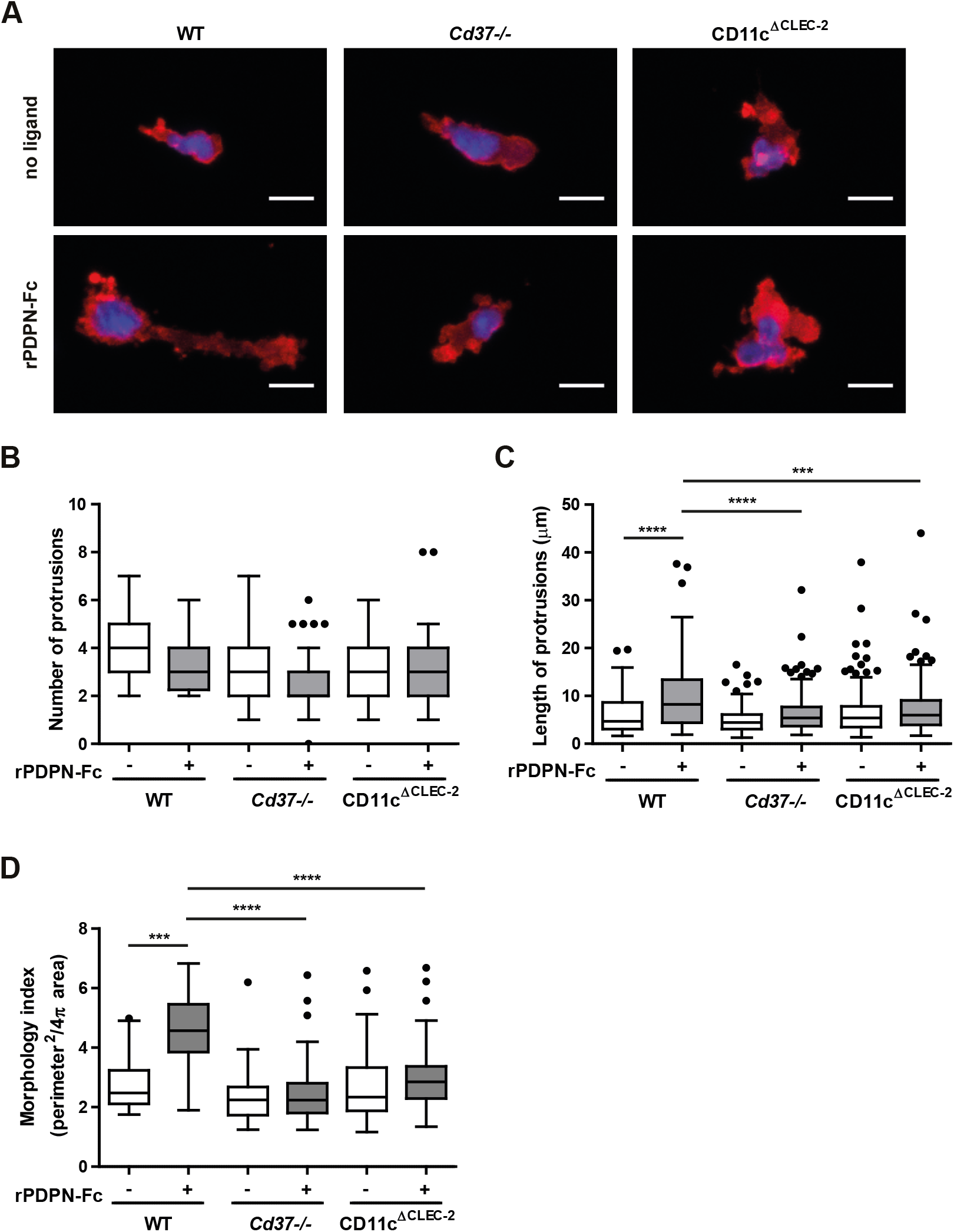
CD37 controls formation of actin protrusions by DCs in response to podoplanin. **(A)** WT (left), *Cd37-/-* (middle) or CD11c^ΔCLEC”2^ (right) DCs were stimulated in a 3D collagen gel with (bottom row) or without (upper row) recombinant podoplanin-Fc (rPDPN-Fc). Cells were stained for F-actin (red) and nucleus (blue) and imaged with a Leica SP5 confocal fluorescence microscope. One representative cell is shown for each condition (overview with more cells is provided in Supplementary Figure 2). Scale bar = 10μm. **(B-D)** Number **(B)** and length **(C)** of actin protrusions, and morphology index **(D)** of WT (left), *Cd37-/-* (middle) or CD11c^ΔCLEC^”^2^ (right) BMDCs upon rPDPN-Fc stimulation (grey boxes) compared to no ligand (white boxes). Data are shown as Tukey Box & whiskers from 3 independent experiments, total n=3 mice per genotype. In Tukey Box & whiskers, black dots are determined as outliers; i.e. data points outside the 25^th^ and 75^th^ percentile, minus or plus the 1.5 interquartile range, respectively. Two-way ANOVA with Tukey’s multiple comparisons, ***p<0.001, ****p<0.0001.

### CLEC-2 recruitment to podoplanin is dependent on CD37

To gain insight into the underlying mechanism by which CD37 controls CLEC-2 response to podoplanin, we analyzed local CLEC-2 recruitment to podoplanin in the presence or absence of CD37 by microcontact printing experiments. Microcontact printing (“stamping”) technology (Van Den Dries et al., 2012; Zuidscherwoude et al., 2017b) enables imaging and analysis of CLEC-2 protein recruitment in the membrane of cells towards recombinant podoplanin protein that is stamped as circular spots (5μm) on glass coverslips. Myeloid cells (RAW264.7 macrophages) were transiently transfected with murine CLEC-2 tagged to GFP (GFP-mCLEC-2) with or without murine CD37 tagged to mCherry (mCD37-mCherry) (Figure 5A and Supplementary Figure 3A), and incubated on coverslips with podoplanin stamps to locally engage CLEC-2 molecules at sites of podoplanin. We determined CLEC-2 membrane expression in the transfected cells to control for differences in transfection efficiency. CLEC-2 membrane expression was comparable between cells transfected with or without mCD37-mCherry (Figure 5B). Cells expressing both GFP-mCLEC-2 and mCD37-mCherry showed a >2-fold higher percentage of cells with local (on-stamp) CLEC-2 enrichment at sites of podoplanin, compared to cells only expressing GFP-mCLEC-2 (Figure 5C-D, Supplementary Figure 3B). These data indicate that CD37 significantly enhances CLEC-2 recruitment to podoplanin.

**Figure 5.**
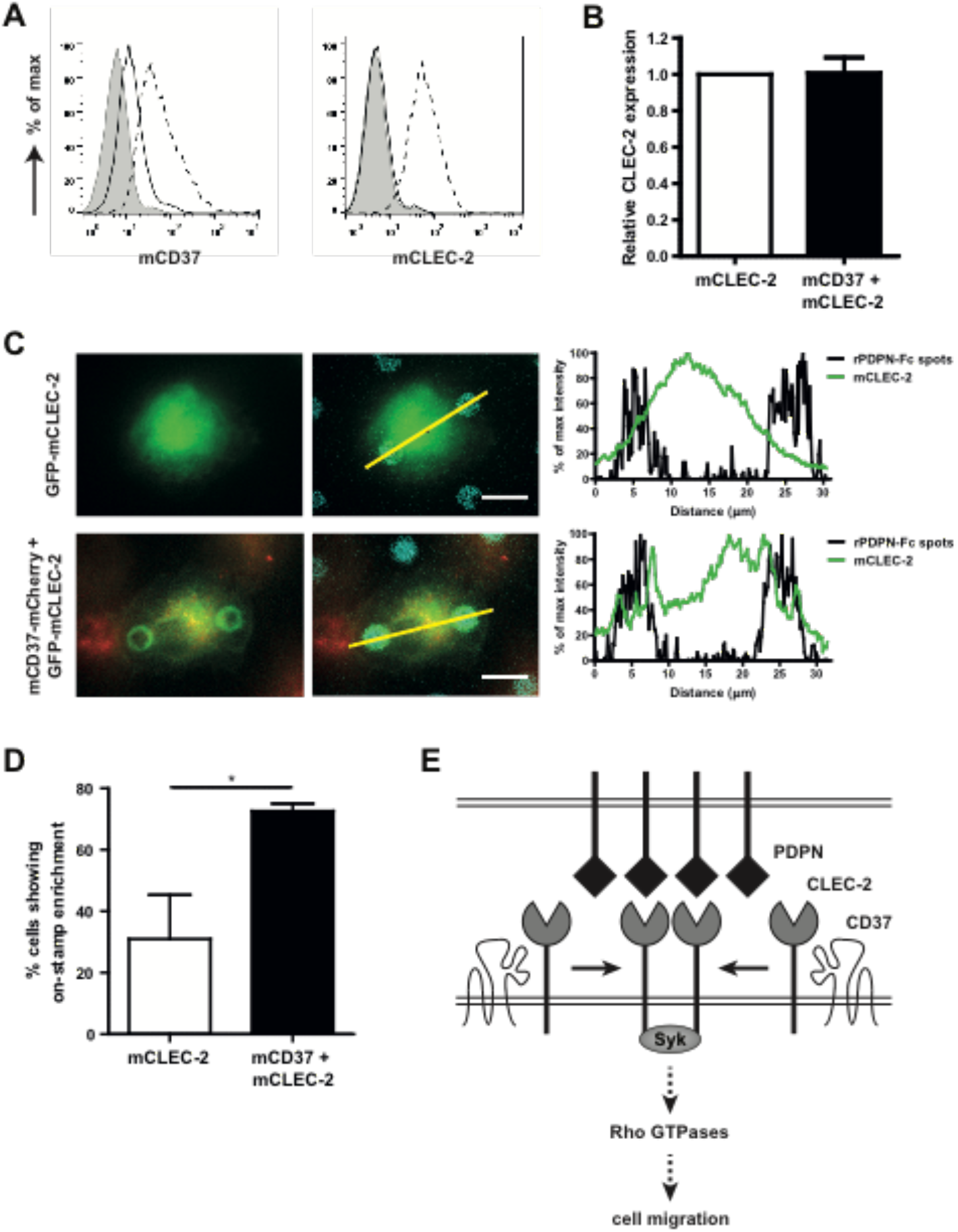
CD37 drives CLEC-2 recruitment to podoplanin and directly interacts with CLEC-2. **(A)** Membrane expression of murine CD37 (mCD37; left) and murine CLEC-2 (mCLEC-2; right) on mock (black line) or transfected (dashed line) RAW macrophages determined by flow cytometry. Histograms show mCD37 (left) or mCLEC-2 (right) membrane expression on live cells that were gated on mCD37-mCherry or GFP-mCLEC-2 positivity, respectively. Grey = isotype control. Raw flow cytometry data are shown in Supplementary Figure 3. **(B)** Relative CLEC-2 membrane expression in RAW cells transfected with GFP-mCLEC-2 (white bar) or mCD37-mCherry (black bar). Flow cytometry results from panel A are normalized per experiment to the level of CLEC-2 membrane expression in RAW cells transfected with GFP-mCLEC-2, which was set at 1. Data are shown as mean+SEM from four independent experiments. **(C)** GFP-mCLEC-2 (upper row) or in combination with mCD37-mCherry (bottom row) transfected RAW macrophages were incubated on recombinant podoplan¡n-Fc (rPDPN-Fc) spots and analyzed after 12 min using an ep¡-fluorescence microscope. Left: Green = GFP, red = mCherry, blue = rPDPN-Fc spots. One representative cell shown per condition (three more representative cells per condition are shown in Supplementary Figure 3). Scale bar = 10μm. Right: Graphs represent intensity profile of GFP-mCLEC-2 (green line) or rPDPN-Fc spot (black line) across the yellow line. **(D)** Percentage of GFP-mCLEC-2 (white bar) or in combination with mCD37-mCherry (black bar) transfected RAW macrophages showing enrichment of GFP-mCLEC-2 on rPDPN-Fc spots. Data are shown as mean+SEM from three independent experiments (in total 54-56 cells per condition). Paired Student’s *t*-test, one-tailed, *p=0.0372. **(E)** Model illustrating that CD37 drives CLEC-2 recruitment in response to its ligand podoplanin and as such regulates DC migration. CLEC-2-induced Syk activation leads to DC migration most likely via changes in the activity of Rho GTPases.

## Discussion

It is well-established that CLEC-2 interaction with podoplanin is essential for DC migration and initiation of the cellular immune response (Acton et al., 2012; Acton et al., 2014; Astarita et al., 2015), still the molecular mechanisms underlying this remain elusive. We identified a novel molecular interaction between CLEC-2 and CD37, which was specific for CD37 as other tetraspanins (CD9, CD63, CD81 and CD151) did not interact with CLEC-2. We discovered that DCs lacking CD37 have decreased CLEC-2 expression at the cell surface, and impaired adhesion, migration velocity and displacement on podoplanin-expressing lymph node stromal cells. Moreover, podoplanin-induced formation of actin protrusions and recruitment of CLEC-2 to podoplanin was impaired in *Cd37-/-* DCs.

For efficient ligand binding and activation of downstream signaling, CLRs have been postulated to depend on spatiotemporal localization into specific microdomains on the plasma membrane (Cambi et al., 2004; Figdor and van Spriel, 2009; Meyer-Wentrup et al., 2007). CLEC-2 has found to be present in clusters on blood platelets (Hughes et al., 2010; Pollitt et al., 2014) and CLEC-2 clusters were reported to be localized in lipid rafts (Manne et al., 2015; Pollitt et al., 2010) by using detergent-resistant membrane extraction. However, this technique also enriches for tetraspanin microdomains (Blank et al., 2007; Claas et al., 2001; Tarrant et al., 2003). We now identified a specific molecular interaction between CLEC-2 and tetraspanin CD37 indicating that CD37 microdomains form the scaffold for CLEC-2 clusters on the plasma membrane of DCs. The finding that CLEC-2 did not interact with other tetraspanins may be explained by recent super-resolution studies of the tetraspanin web in which individual tetraspanins were found in separate nanoclusters (100-120nm) at the cell surface of B cells and DCs (Zuidscherwoude et al., 2015). Our data are in line with the demonstration that expression and stabilization of the CLEC-2 homologous family member Dectin-1 at the plasma membrane of macrophages was dependent on CD37 (Meyer-Wentrup et al., 2007). The role of CD37 in stabilizing CLRs on the plasma membrane may also underlie our finding that CLEC-2 surface expression is decreased on *Cd37-/-* splenocytes compared to WT splenocytes, indicating that CLEC-2 turnover is increased in absence of CD37.

Our results show that CLEC-2+ *Cd37-/-* DCs have an impaired capacity to adhere to podoplanin-expressing LECs, and demonstrate lower migration velocity and displacement. This is in accordance with studies showing impaired migration of *Cd37-/-* DCs *in vivo* (Gartlan et al., 2013; Jones et al., 2016). In DC biology, tetraspanin interactions with adhesion receptors or MHC molecules are particularly important for antigen presentation and T cell activation (Gartlan et al., 2010; Jones et al., 2016; Rocha-Perugini et al., 2017; Sheng et al., 2009). Now, we demonstrate a specific role for CD37 in controlling CLEC-2 function in migration of DCs. *Cd37-/-* DCs show impaired actin protrusion formation upon podoplanin stimulation, which is highly similar to the phenotype of CD11c^ΔCLEC-2^ DCs. Rearrangements of the actin cytoskeleton and subsequent cell movement are controlled by Rho GTPases, including RhoA and Rac1. RhoA increases actomyosin contractility via its interaction with Rho kinases (Parri and Chiarugi, 2010), whereas Rac1 supports actin polymerization, spreading of lamellipodia and formation of membrane ruffles (Nobes and Hall, 1995; Olson and Sahai, 2009). Since activity of Rho GTPases has been shown to change upon CLEC-2 activation by podoplanin or rhodocytin (i.e. downregulation of RhoA and upregulation of Rac1) (Acton et al., 2012), we postulate that the underlying molecular mechanism of the defective cell migration of *Cd37-/-* DCs is due to deregulation of intracellular Rho GTPases as a consequence of impaired recruitment of CLEC-2 to its ligand podoplanin. This is supported by a recent study demonstrating impaired activation of Rac-1 in toxin-activated adherent *Cd37-/-* bone marrow-derived DCs (Jones et al., 2016). Altogether, these data support a model in which CD37 is important for CLEC-2 recruitment in the plasma membrane of myeloid cells upon podoplanin binding, which results in Syk activation and changes in Rho GTPase activity (e.g. increased Rac1 and decreased RhoA activation) leading to cell migration (Figure 5E).

Besides activation of Rho GTPases, CLRs can initiate intracellular Syk-dependent signaling cascades that induce cytokine production (Mócsai et al., 2010). We found that *Cd37-/-* myeloid cells produce higher IL-6 levels compared to WT cells upon stimulation with the CLEC-2 ligand rhodocytin, which is in line with previous reports showing production of pro-inflammatory cytokines (i.e. IL-6 and TNFα) by neutrophils and RAW macrophages upon stimulation with rhodocytin (Chang et al., 2010; Kerrigan et al., 2009). Increased IL-6 expression upon CLR stimulation has also been shown in *Cd37-/-* macrophages upon stimulation with the Dectin-1 ligand curdlan (Meyer-Wentrup et al., 2007). Additionally, IL-10 production by RAW macrophages and BMDCs co-stimulated with LPS and anti-CLEC-2 Fab fragments could be reversed by Syk inhibition (Mourão-Sá et al., 2011). Our results suggest that CD37 directly controls Syk signaling downstream of hemITAM CLRs and as such inhibits cytokine production, probably by stimulating phosphatase activity (Carmo and Wright, 1995; Chattopadhyay et al., 2003; Wright et al., 2004). Syk activation has been reported to be negatively regulated by SH2 domain-containing protein tyrosine phosphatase 1 (SHP1) (Mócsai et al., 2010; Zhang et al., 2000). SHP1 and CD37 have been shown to associate via the N-terminal ITIM-like domain of CD37 in chronic lymphocytic leukemia cells, which induced tumor cell death via negative regulation of AKT-mediated pro-apoptotic signaling (Lapalombella et al., 2012). The binding of cytoplasmic signaling proteins, like protein kinase C (PKC) (Zhang et al., 2001; Zuidscherwoude et al., 2017b), Rac1 (Tejera et al., 2013), and suppressor of cytokine signaling 3 (SOCS3) (de Winde et al., 2016) have been reported for different tetraspanins (also reviewed in (van Deventer et al., 2017)).

In conclusion, our results demonstrate that CD37 is required for ligand-induced CLEC-2 responses via a direct molecular interaction leading to immune cell activation and DC migration. Furthermore, this study supports a general mechanism of tetraspanin-mediated membrane organization and movement of CLRs in the plasma membrane, which underlies cytoskeletal changes and cell migration.

## Materials and methods

### Mice

*Cd37-/-* mice (male, average age of three months) were generated by homologous recombination (Knobeloch et al., 2000) and fully backcrossed to the C57BL/6J background (van Spriel et al., 2004). *Cd37+/+* (WT) littermates were matched for age and gender. *Cd37+/+ and Cd37-/-* mice were bred in the Central Animal Laboratory of Radboud University Medical Center. CD11c^ΔCLEC-2^ mice (C57BL/6J background), selectively lacking CLEC-2 in CD11c+ cells, were generated by crossing Cd11c-cre and Clec1bfl/fl mice as previously described (Acton et al., 2014). All murine studies complied with European legislation (directive 2010/63/EU of the European Commission) and were approved by local authorities (CCD, The Hague, the Netherlands, and Institutional Animal Ethics Committee Review Board, Cancer Research UK and the UK Home Office, United Kingdom) for the care and use of animals with related codes of practice. All mice were housed in top-filter cages and fed a standard diet with freely available water and food.

### Isolation of whole blood

Mice were anesthetized with isoflurane and whole blood was harvested via retro-orbital punction and collected in a tube containing acid-citrate-dextrose mixture (ACD; 25g/L sodium citrate, 20g/L glucose (both from Sigma-Aldrich, Zwijndrecht, The Netherlands) and 15g/L citric acid (Merck, Amsterdam, The Netherlands) to prevent clotting.

### Cell/culture/

Bone marrow-derived DCs (BMDCs) were generated by culturing total murine bone marrow cells in complete medium (RPMI 1640 medium (Gibco, via Thermo Fisher Scientific, Bleiswijk, The Netherlands), 10% fetal calf serum (FCS; Greiner Bio-One, Alphen a/d Rijn, The Netherlands), 1% UltraGlutamine-I (UG; Lonza, Breda, The Netherlands), 1% antibiotic-antimycotic (AA; Gibco, via Thermo Fisher Scientific, Bleiswijk, The Netherlands) and 50μM β-mercapto-ethanol (Sigma-Aldrich, Zwijndrecht, The Netherlands), containing 20ng/mL murine granulocyte-macrophage colony stimulating factor (mGM-CSF; Peprotech, via Bio-Connect, Huissen, The Netherlands), as adapted from previously described protocols (Lutz et al., 1999). On day 6, BMDCs were additionally stimulated with 10ng/mL LPS (Sigma-Aldrich, Zwijndrecht, The Netherlands) for 24h, unless stated otherwise.

Primary human dermal lymphatic endothelial cells (LEC) were purchased from PromoCell and cultured in manufacturer’s recommended medium (Endothelial Cell Growth Medium MV2, PromoCell, Heidelberg, Germany) supplemented with 35μg/mL gentamicin (Gibco, via Fisher Scientific - UK Ltd, Loughborough, UK). As previously described (Ahmed et al., 2011), LECs were dissociated using a 2:1 ratio of trypsin (2.5mg/ml) to EDTA (0.02%) (both from Sigma-Aldrich, Paisley, UK) and seeded on 12-well tissue culture plates coated with 2% gelatin (Sigma-Aldrich, Paisley, UK). Seeding density was chosen to yield a confluent LEC monolayer within 24h. TNF-alpha (TNFα; 100U/ml; R&D Systems, via Bio-Techne Ltd, Abingdon, UK) was added to confluent LEC monolayers for 24h before analyzing static adhesion and migration of DCs (Johnson and Jackson, 2013; Maddaluno et al., 2009; Podgrabinska et al., 2009).

RAW264.7 murine macrophages (RAW; originally from ATCC) were cultured in RPMI 1640 medium (Gibco, via Thermo Fisher Scientific, Bleiswijk, The Netherlands) supplemented with 10% FCS (Greiner Bio-One, Alphen a/d Rijn, The Netherlands), 1% UG (Lonza, Breda, The Netherlands) and 1% AA (Gibco, via Thermo Fisher Scientific, Bleiswijk, The Netherlands). The human embryonic kidney (HEK)-293T (HEK-293 cells expressing the large T-antigen of simian virus 40) cell line was cultured in complete DMEM medium (Sigma-Aldrich, Zwijndrecht, The Netherlands) containing 10% fetal calf serum (Gibco, via Thermo Fisher Scientific, Loughborough, UK), 4mM L-glutamine, 100U/ml penicillin and 100μg/ml streptomycin (Thermo Fisher Scientific, Loughborough, UK).

### Constructs and transfection

mCD37-pmCherry was generated by fluorescent protein swap of GFP from mCD37-pEGFP (Meyer-Wentrup et al., 2007) with mCherry from pmCherry-N1 (Clontech, via Takara Bio Europe, Saint-Germain-en-Laye, France) using AgeI and BsrGI restriction sites (New England Biolabs (via Bioké, Leiden, The Netherlands)). RAW macrophages (5x10^5^ cells/transfection) were transfected with 0.5μg mCD37-mCherry and/or 0.5μg pAcGFP-mCLEC-2 as previously described (Pollitt et al., 2014) using FuGENE HD according to manufacturer’s instructions (Promega, Leiden, The Netherlands).

Human (h)CLEC-2-MYC construct with C-terminal MYC tags (Fuller et al., 2007) was generated in the pEF6 expression vector (Invitrogen, via Thermo Fisher Scientific, Loughborough, UK). The FLAG-human CD37 and other human tetraspanin constructs were produced using the pEF6 expression vector with an N-terminal FLAG tag as described previously (Protty et al., 2009). HEK-293T cells were transiently transfected with hCLEC-2-MYC and FLAG epitope-tagged human tetraspanin expression constructs using polyethylenimine (Sigma-Aldrich, Paisley, UK) as described previously (Ehrhardt et al., 2006; Noy et al., 2015).

### Immunoprecipitation

Transfected HEK-293T cells were lysed in 1% digitonin (Acros Organics (via Thermo Fisher Scientific, Loughborough, UK) and immunoprecipitated with anti-FLAG antibody (clone M2; Sigma-Aldrich, Paisley, UK) as described previously (Haining et al., 2012). Western blots were stained with anti-MYC (clone 9B11; Cell Signaling Technology, via New England Biolabs Ltd, Hitchin, UK) or rabbit anti-FLAG (Sigma-Aldrich, Paisley, UK) antibodies, followed by IRDye^®^ 680RD- or 800CW-conjugated secondary antibodies (LI-COR Biotechnology, Cambridge, UK), and imaged using the Odyssey Infrared Imaging System (LI-COR Biotechnology, Cambridge, UK).

### Flow cytometry

Murine whole blood was incubated in phosphate-buffered saline (PBS; Braun, Aschaffenburg, Germany) in presence of 1mM EDTA (Amresco, via VWR International, Amsterdam, The Netherlands) to prevent platelet aggregation, and 2% normal goat serum (NGS; Sigma-Aldrich, Zwijndrecht, The Netherlands) to block Fc receptors for 15 min at 4°C. Next, murine blood cells were stained with anti-mouse CLEC-2 (clone INU1; a kind gift from Bernhard Nieswandt, University of Würzburg, Germany) or appropriate isotype control, and subsequently with goat-anti-rat IgG-APC (BD Biosciences, Vianen, The Netherlands). To discriminate blood platelets, staining with anti-mouse CD41-PE (clone MWReg30; BD Biosciences, Vianen, The Netherlands) was performed.

Splenocytes or cell lines were stained for 30 min at 4°C in PBS (Braun, Aschaffenburg, Germany) containing 1% bovine serum albumin (BSA; Roche, Almere, The Netherlands) and 0.05% NaN_3_, and supplemented with 2% NGS (Sigma-Aldrich, Zwijndrecht, The Netherlands), with the following primary anti-mouse antibodies: CLEC-2 (clone INU1; a kind gift from Bernhard Nieswandt, University of Würzburg, Germany), CD37 (clone Duno85; Biolegend, London, UK), B220-FITC (CD45R, clone RA3-6B2; Biolegend, London, UK), CD11c-Alexa488 (clone N418; Biolegend, London, UK), NK1.1-PE (clone PK136; BD Biosciences, Vianen, The Netherlands), GR1-PE (clone RB6-8C5; Biolegend, London, UK), CD11b-PerCP (clone M1/70; Biolegend, London, UK), CD3-biotin (CD3ε, clone 145-2C11; eBioscience, via Thermo Fisher Scientific, Bleiswijk, The Netherlands), or appropriate isotype controls. This was followed by incubation with streptavidin-PerCP (Biolegend, London, UK) or goat-anti-rat IgG Alexa647 (Invitrogen, via Thermo Fisher Scientific, Bleiswijk, The Netherlands). Stained cells were analyzed using FACSCalibur (BD Biosciences, Vianen, The Netherlands) or CyanADP (Beckman Coulter, Woerden, The Netherlands) flow cytometer, and FlowJo software (TreeStar Inc., Ashland, OR, USA).

### Rhodocytin stimulation and cytokine production

Single cell suspensions of splenocytes were generated by passing spleens through a 100μm cell strainer (Falcon, via Corning, Amsterdam, The Netherlands). To lyse erythrocytes, splenocytes were treated with ACK lysis buffer (0.15M NH_4_Cl, 10mM KHCO_3_ (both from Merck, Amsterdam, The Netherlands), 0.1mM EDTA (Sigma-Aldrich, Zwijndrecht, The Netherlands); pH 7.3) for 2 min on ice. Splenocytes (5x10^5^ cells) were stimulated with either 15μg rhodocytin (purified and kindly provided by Prof. Johannes Eble as previously described (Eble et al., 2001)), PMA/ionomycin (100ng/mL and 500ng/mL, both from Sigma-Aldrich, Zwijndrecht, The Netherlands), or with complete RPMI 1640 medium (unstimulated), and incubated overnight at 37°C, 5% CO_2_. To measure IL-6 levels in the supernatant of stimulated cell cultures, NUNC Maxisorp 96-well plates (eBioscience, via Thermo Fisher Scientific, Bleiswijk, The Netherlands) were coated with capture anti-mouse IL-6 antibody (MP5-20F3, 2μg/mL; BD Biosciences, Vianen, The Netherlands) in 0.1M carbonate buffer (pH 9.6) overnight at 4°C. Wells were blocked with PBS (Braun, Aschaffenburg, Germany) containing 1% BSA (Roche, Almere, The Netherlands) and 1% FCS (Greiner Bio-One, Alphen a/d Rijn, The Netherlands) for 1 hour at room temperature (RT), washed, and incubated with 50μL of sample and standard (2-fold serial dilutions starting from 10000pg/ml) (eBioscience, via Thermo Fisher Scientific, Bleiswijk, The Netherlands). After 2-hour incubation at RT, wells were incubated with biotinylated anti-mouse IL-6 (MP5-32C11, 1μg/ml, BD Biosciences, Vianen, The Netherlands) for 1 hour at RT, followed by incubation with HRP-conjugated streptavidin (1:5000, Invitrogen, via Thermo Fisher Scientific, Bleiswijk, The Netherlands) for 30 min at RT. Complexes were visualized using TMB substrate (Sigma-Aldrich, Zwijndrecht, The Netherlands), and reactions were stopped by adding 0.8M H_2_SO_4_ (Merck Millipore, Amsterdam, The Netherlands). Absorbance was measured at 450nm using an iMark plate reader (Bio-Rad, Veenendaal, The Netherlands).

### Static adhesion and migration assay

Adhesion was assessed by direct microscopic observation as previously described (Butler et al., 2005). LEC in 12-well plates were washed three times with Medium 199 supplemented with 0.15% BSA Fraction V 7.5% (M199BSA, both from Gibco, via Thermo Fisher Scientific, Loughborough, UK) to remove residual cytokines. BMDCs (1x10^6^) were added on top of the LEC monolayer and incubated for 10 min at 37°C, 5% CO_2_. Non-adherent cells were removed from the LECs by gentle washing three times with M199BSA medium. Imaging was performed using an Olympus Invert X70 microscope enclosed at 37°C. Digital recordings of five fields of view of the LEC surface were made using phase-contrast microscopy immediately and 10 min after washing away non-adherent cells. In between, time-lapse imaging (1 image every 10 sec) was performed for 5 min of one field of view to assess cell migration at 37°C. Digitized recordings were analyzed off-line using Image-Pro software (version 6.2, DataCell, Finchampstead, UK). The numbers of adherent cells were counted in the video fields, averaged, converted to cells per mm^2^ using the calibrated microscope field dimensions, and multiplied by the known surface area of the well to calculate the total number of adherent cells. This number was divided by the known total number of cells added to obtain the percentage of the cells that had adhered. Cell tracks of live cells were analyzed using the Manual Tracking plugin in Fiji/ImageJ software. Migration velocity was calculated as the length of each cell path/time (μm/min), and the *xy* trajectories were converted into mean square displacement (MSD, in μm^2^) as previously described (van Rijn et al., 2016).

### 3D protrusion assay

BMDCs (0.3x10^6^) were seeded into a 3D collagen (type I, rat tail)/matrigel matrix (both from Corning, via Thermo Fisher Scientific, Loughborough, UK) supplemented with 10% minimum essential medium alpha medium (MEMalpha, Invitrogen, via Thermo Fisher Scientific, Loughborough, UK) and 10% FCS (Greiner Bio-One, Stonehouse, UK) on glass-bottomed cell culture plates (MatTek Corporation, Bratislava, Slovakia). For CLEC-2 activation, 20μg/mL recombinant podoplanin-Fc (rPDPN-Fc; Sino Biological Inc., Beijing, China) (Acton et al., 2012) was added to the gel as all components were mixed. Gels were incubated overnight at 37°C, 5% CO_2_. The next day, cultures were fixed with 4% paraformaldehyde (PFA; Merck, Nottingham, UK) for 3h at RT, followed by blocking and permeabilization with 3% BSA (Roche, West Sussex, UK), 1% normal mouse serum (NMS) and 0.3% Triton X-100 in 0.1M Tris (all from Sigma-Aldrich, Paisley, UK) for 2h at RT before staining. F-actin and cell nuclei were visualized using TRITC-phalloidin and DAPI, respectively (1:1000 dilution, both from Invitrogen, via Thermo Fisher Scientific, Loughborough, UK). Cells were imaged on a Leica SP5 confocal microscope, and analyzed using Fiji/ImageJ software. Z stacks of 120μm (10μm/step) were projected with ImageJ Z Project (maximum projection), and number and length of protrusions were measured. Cell morphology (= perimeter2/4πarea) was calculated using the area and perimeter of cells by manually drawing around the cell shape using F-actin staining.

### Microcontact printing

PDMS (poly(dimethylsiloxane)) stamps with a regular pattern of 5μm circular spots were prepared as previously described (Van Den Dries et al., 2012). Stamps were incubated with 200μg/mL rPDPN-Fc (Sino Biological Inc., Beijing, China) for CLEC-2 stimulation, and 10μg/mL donkey anti-rabbit IgG (H&L)-Alexa647 (Invitrogen, via Thermo Fisher Scientific, Loughborough, UK) to visualize the spots, diluted in PBS (Braun, Aschaffenburg, Germany) for 1 hour at RT. Stamps were washed with demineralized water and dried under a nitrogen stream. The stamp was applied to a cleaned glass coverslip for 1 min and subsequently removed. Transfected RAW macrophages were seeded on the stamped area and incubated for 12 min at 37°C. Cells were fixed with 4% PFA (Merck, Darmstadt, Germany) for 20 min at RT. Samples were washed with PBS (Braun, Aschaffenburg, Germany) and demineralized water (MilliQ; Merck Millipore, Amsterdam, The Netherlands) and embedded in Mowiol (Sigma-Aldrich, Zwijndrecht, The Netherlands). Imaging was performed on an epi-fluorescence Leica DMI6000 microscope, and plot profiles were created using Fiji/ImageJ software. For the population of cells in contact with rPDPN-Fc spots, we determined the percentage showing on-stamp enrichment of mCLEC-2-GFP by independent visual analysis with support of Fiji/ImageJ software.

### Statistics

Statistical differences between two groups (e.g. WT and *Cd37-/-* cells) regarding IL-6 production, adhesion, CLEC-2 enrichment and NFAT-luciferase activity were determined using (un)paired Student’s *t*-test or non-parametric Mann-Whitney test (in case of non-Gaussian distribution). Statistical differences between three groups (e.g. WT, *Cd37-/-* and CD11c^ΔCLEC-2^ cells) or two or more parameters (e.g. genotype and time) were determined using two-way ANOVA with Sidak’s or Tukey’s multiple comparisons test. Statistical tests were performed using GraphPad Prism software, and all differences were considered to be statistically significant at p≤0.05.

## Acknowledgements

We thank Prof. Steve Watson (University of Birmingham, UK; British Heart Foundation Chair (CH03/03)) and Dr. Mark Wright (Monash University, Melbourne, Australia) for discussion and critical reading of the manuscript. We thank Jing Yang for her contribution to the research during her function as research assistant in the lab of Dr M.G. Tomlinson. C.M. de Winde was supported by an Erasmus+ Staff Mobility Grant, and was awarded a NWO Rubicon Postdoctoral Fellowship (019.162LW.004). A.L. Matthews was supported by a Biotechnology and Biological Sciences Research Council PhD Studentship. N.D. Tomlinson was supported by a Medical Research Council PhD Studentship. J.A.E. isolates rhodocytin within a project financed by the Deutsche Forschungsgemeinschaft (grant: SFB1009 A09). H.M. McGettrick was supported by an Arthritis Research UK Career Development Fellowship (19899). C.G. Figdor is recipient of a NWO Spinoza award, a European Research Council Advanced Grant PATHFINDER (269019), and a Koningin Wilhelmina Onderzoeksprijs award (KUN2009-4402) from the Dutch Cancer Society. S.E. Acton is recipient of a Cancer Research UK Career Development Fellowship (CRUK-A19763) and is supported by the Medical Research Council (MC_U12266B). M.G. Tomlinson was supported by a Medical Research Council New Investigator Award (G0400247) and a British Heart Foundation Senior Fellowship (FS/08/062/25797), the latter of which also supported J. Yang. A.B. van Spriel is recipient of a Netherlands Organization for Scientific Research Grant (NWO-ALW VIDI grant 864.11.006), a NWO Gravitation Programme 2013 grant (ICI-024.002.009), a Dutch Cancer Society Grant (KUN2014-6845), and was awarded a European Research Council Consolidator Grant (Secret Surface, 724281). The authors declare no competing financial interests.

